# Protein structure and function analyses to understand the implication of mutually exclusive splicing

**DOI:** 10.1101/292813

**Authors:** Su Datt Lam, Christine Orengo, Jonathan Lees

## Abstract

Alternative splicing (AS) has been suggested as one of the major processes expanding the diversity of proteomes in multicellular organisms. Mutually exclusive exons (MXE) provide one form of AS that is less likely to disrupt protein structure and is over-represented in the proteome compared to other forms of AS. We used domain structure information from the CATH classification to perform a systematic structural analysis of the effects of MXE splicing in high quality animal genomes (e.g. human, fly, mouse and 2 fishes) and we were able to annotate approximately 50% of MXE events with structural information. For those MXE events which can be mapped to a structure, we found that although embedded in domains, they were strongly enriched in surface exposed residues. We also demonstrated that the variable residues between splicing events lie close to known and/or predicted functional sites. We present some examples of MXE events in proteins that have important roles in cells. This work presents the first large scale systematic study of the structural/functional effects of MXE splicing using predominantly domain based modelling and functional annotation tools. Our study supports and expands on previous work in this field and helps to build a picture of how MXE events facilitate evolution of new functions.

## INTRODUCTION

Alternative splicing (AS) has been proposed as an important process for expanding the diversity of proteomes in multicellular organisms (1, 2). It refers to the assemblage and rearrangement of different exons and introns of a gene during pre-mRNA splicing such that different mRNAs and thus proteins are produced from the same gene. AS is common in humans and other animals producing different isoforms in different developmental stages, tissues or disease states) (3). It has been linked to various diseases and cancers (4, 5) and can lead to the tissue-specific rewiring of protein-protein networks (6). Light and Elofsson showed that 36% of the isoforms in Swiss-Prot alter the domain architecture of the proteins (7). Liu and Altman identified a total of 24 protein domains that more commonly undergo AS than other proteins in human (8). The most prevalent domains were the repeating cadherin domain and domains that contain intrinsically disordered regions (6, 8). Cadherin is a type of trans-membrane protein which is predominantly involved in the process of cell communication/signalling, development and apoptosis. Intrinsically disordered regions are prominently found among the hubs in a protein-protein interaction network and it is believed that changes in these proteins are important for mediating the remodelling of protein-protein interaction networks (9, 10).

A recent large-scale RNA-seq experiment identified putative splice variants for 72% of the annotated human genes (11). Proteins produced by AS isoforms may have modified structures and functions. Tress et al demonstrated that AS can lead to wide range of structural outcomes, although a large number of these were difficult to rationalise as functional structures (12). It was found that up 60% of the AS isoforms that could be modelled by Tress and co-workers, lacked a major part of a domain (12), indicating that the AS variants may well not produce functional proteins.

Furthermore, although AS events have been shown to be abundant at the transcript level (11, 13), recent mass spectrometry (MS) analysis showed a much lower prevalence of AS at the protein level (14, 15). However various technical limitations of proteomics and the low splicing abundance of many AS variants could mean that many protein isoforms that are not detected could be identified using more targeted experiments. In addition, ribosome profiling has been employed to detect AS at the protein level (16, 17). Although Weatheritt et al. demonstrated that at least 75% of human exon-skipping events were associated with ribosomes (17), there has been some controversy regarding these observations (18). Therefore, many pieces of evidence point to a large proportion of alternative splicing events arising from noise in the transcriptional machinery (19).

However, although the splice isoforms identified in proteomics experiments have low coverage, they have been found to be highly enriched with mutually exclusive exons (MXE) both in human and mouse analyses (14). MXEs are characterised by the splicing of exons in a coordinated manner such that two or more splicing events are not independent. Only one out of the two exons is retained, while the other one is spliced out (Figure 1). It is generally believed that MXEs originated from exon duplication, and often show high sequence similarity. They are usually similar in the length (typically the number of residues in the alpha-helical and beta-strands regions are conserved, but there is variation in the lengths of the loop regions), conserved splice site patterns (e.g. GT-AG, GC-AG, and AT-AC), same reading frame and high sequence identity (20, 21). MXE events have been shown to be highly conserved throughout vertebrates (22, 23).

**Figure 1.**
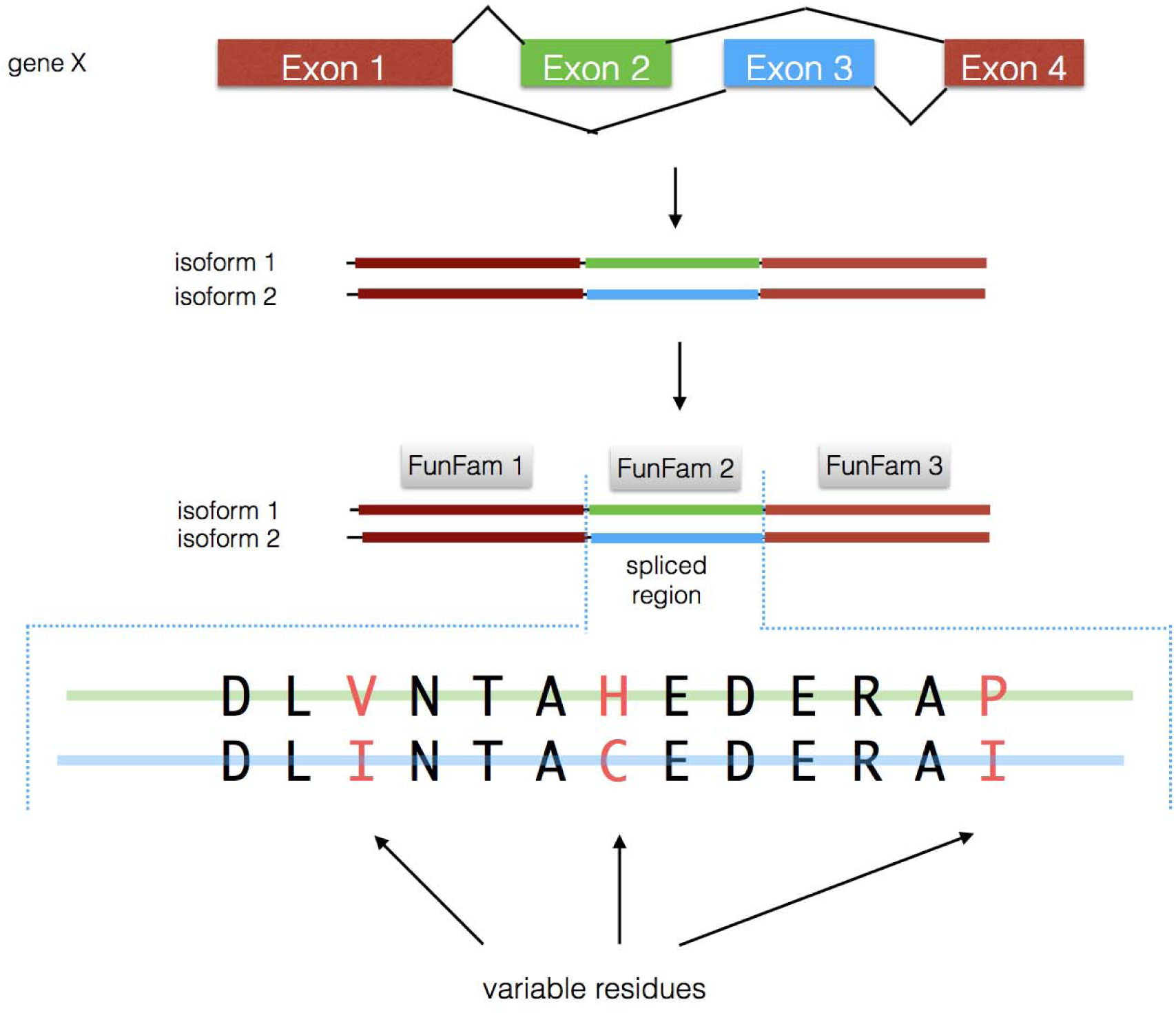
Identifying the variable residues between splice events. Isoforms with evidence of a MXE event are aligned to FunFams and then differences between amino acids are identified from the aligned residues. Note: For the cases which are not mapped to a FunFam, the isoforms are aligned to a structure in the PDB whose sequence best matches the isoforms.

An analysis of MXE events, to measure the structural and functional effects of AS showed that only 1 out of the 60 human MXE events disrupted a Pfam domain (measured by loss or gain of five or more residues) (14). Abascal and co-workers managed to directly map 9 of the 60 human MXE to known PDB structures. However, this study only used known structure and did not expand to analysis of structural models. From these and other studies it is clear that MXE splice events have relatively subtle effects on protein folds compared to other forms of AS such as exon skipping. Recent analysis also demonstrated human MXEs to be embedded within folded structural domains (23). The combination of these two observations means that MXE events would be well suited to a structural modelling approach.

As with the structural analyses, analysis of functional consequences of MXE events have been limited to small scale studies. Examples have been found where the MXE splicing modulates the binding of the protein to another substrate (protein, ion) (15, 22) or disturbs the catalytic site or an allosteric site. MXEs have been proven to be have other biological effects, such as regulation of mammalian expression level (24) and voltage dependence of ion channels (25).

As discussed above there is much more evidence for MXEs to be of functional importance compared to other AS events. The mechanism of MXEs (i.e. substituting a similar length sequence region) should generally lead to preservation of the overall structure and therefore allow us to more easily identify the consequences of the sequence changes brought about by the AS event at a structural level. However, to date, structural modelling has all been on a small scale and no systematic study has been carried out.

To explore the structural and functional consequences of MXE splicing events in five species (human, fly, mouse, fugu fish and zebra fish), in a systematic way, we mapped the isoforms to functional families (FunFams) in the CATH domain structure classification (26). Relatives in FunFams have highly similar structures and functions (27, 28). Splice regions were mapped to known structures where available, or homology models built using the in-house FunMod modelling pipeline (29, 30). If no model could be built, splice events were aligned to known structures using an in-house HMM-based protocol.

We used the McLachlan physiochemical matrix (31) to quantify the amino acid changes of variable residues identified for a given MXE event. We determined whether the variable residues in an MXE event are exposed to solvent and in the vicinity of any known functional sites e.g. catalytic residues (from the Catalytic Site Atlas (CSA) (32)), protein-protein interaction sites, protein-small molecule interaction sites (from the Inferred Biomolecular Interaction Server (IBIS) (33)), allosteric sites (predicted using the A-SITE predictor) and FunSites (in-house predictions, based on conserved sites in FunFam alignments and proven to be enriched with known functional sites (27, 28)). We also mapped residues mutated in cancer, from the Catalogue of Somatic Mutations in Cancer database (COSMIC) (34) to MXE events to determine how these mutations relate to MXE events.

Our analysis shows that a large scale computational analysis of structural/functional impacts of MXE events is possible using a homology modelling platform to provide predicted structural data for these events. The study exploits this structural data to detect a significant proportion of MXE events from multiple species that are likely to have important effects on protein function. Notably, MXE splicing consistently alters surface residues and we can demonstrate that many of these events are close to known or predicted functional sites in the proteins and therefore likely to produce functional changes.

## MATERIAL AND METHODS

### Identifying the amino acid sequence region involved in MXE events

We downloaded the annotations of predicted MXEs for *Drosophila melanogaster* from the Kassiopeia resource (21) which corresponded to FlyBase 5.36 (35) and kept the MXE events and associated protein sequences that we could map to this genome. For example, we did not make use of predicted exons not in the core genome annotation and only used examples where the translations from chromosome sequence exactly matched the FlyBase translation.

The Kassiopeia resource does not provide MXE download annotations for the other four species (human, mouse, zebra fish and fugu fish). In order to identify MXEs for these species, we did the following. Based on Ensembl version 87, we obtained the mRNA transcripts and identified sets of genes with MXEs. We selected the longest transcript as our reference and compared all the other transcripts against the reference to look for MXE exons. We made sure that the exons were consecutive in the DNA sequence, but never occurred together in the same transcript, were similar in length and had the same reading frame. After that, we translated all the MXE exons into amino acid sequences and compared the sequences against each other with BLAST (36), setting an e-value threshold of 0.005 and minimum sequence identity 25%. Because we wanted to be more confident of the homology we filtered out instances where either MXE was < 8 residues. In order to help with our structural studies, we also filtered out those with problematic length differences by checking that the absolute value of the log ratio of the MXE sequence lengths was > 0.25.

### Predicting the function of MXE genes

We used version 6.8 of the DAVID function annotation tool suite (37), last updated in October 2016, to identify functional enrichments of identified MXE genes. DAVID first maps the gene list (e.g. FlyBase gene id or Ensembl gene id) to over 40 different types of biological annotations (e.g. UniProt sequence feature, Enzyme Commission term, Gene Ontology terms etc.) and outputs the significantly most enriched biological annotations. We used the whole genome background for the species, respectively.

### Annotating MXE events with structural information

We mapped the splice isoforms to CATH functional families (FunFams). FunFams are a sub-classification of CATH protein domain superfamilies (26) that cluster relatives likely to have very similar structures and functions. They are generated using the FunFHMMer protocol (27) that detects similarities in sequence patterns (highly conserved positions and specificity determining positions). Positions that are highly conserved across a superfamily are generally important for the stability, folding or function of the protein domain. Specificity-determining positions are positions that are conserved within and unique to a particular cluster, responsible for a specific function and usually involved in functional divergence from other clusters. Therefore, the FunFam protocol uses agglomerative clustering to iteratively merge clusters of relatives but clusters having different specificity determining residues are not merged (27).

The domains within a given FunFam have been demonstrated to be structurally coherent (28, 29). The functional purity of the FunFams has been demonstrated by validating against experimentally determined proteins from the Enzyme Commission and also by checking whether known functional sites coincide with highly conserved residues in the Multiple Sequence Alignments (MSAs) of FunFams (27, 28). CATH FunFams have been shown to be more functionally pure than Pfam domain families (28). Functional predictions based on FunFams were ranked amongst the top five methods for the "Molecular Function" category and the "Biological Process" category in the most recent CAFA International Function Prediction experiments (CAFA2, 39, CAFA3, P. Radivojac and I. Friedberg, personal communication).

All the splice events were scanned against the library of CATH v4.1 FunFam HMMs (39) using HMMER (40). DomainFinder3 (41) was used to determine which CATH-Gene3D FunFams they belonged to. We only considered matches with a HMMER E-value of less than 0.001. For FunFams that had a known domain structure, we annotated the splice events using the FlyBase/Ensembl to PDB mapping. For FunFams that had no relative of known structure, we used the FunMod modelling pipeline (29, 30) which exploits the MODELLER (42) algorithm to build structural models. We used normalised DOPE (43) and GA341 (44) to assess the quality of the models. Only good quality models with a negative normalised DOPE score and a GA341 score of more than 0.7 were included in this analysis. For those splice events where we failed to build a model, we mapped them to the structural representative of the respective FunFam. To ensure that we chose a structural representative that represents the FunFam well, the structural domain with the highest cumulative SSAP structural similarity score and the best X-ray resolution was used. SSAP is a well-established structural comparison method (45). The SSAP score ranges from 0 to 100, with a score of 100 for identical structures.

The two splice isoforms for an MXE event were aligned to other relatives in the FunFam using the multiple sequence alignment tool MAFFT (46) and the function mafft-add. We extracted the alignment between the structural representative of the FunFam and the two splice isoforms. Based on this alignment, we extracted the variable residues between the two isoforms (See Figure 1).

For those MXE events which are not included in FunFams, we scanned the isoforms against the libraries of HMMs built from non-redundant structures in various resources, CATH v4.1 HMMs (i.e. including non-FunFam CATH domains), SCOPe 2.06 HMMs (47) and PDB70 June 2017 HMMs (48) using either HMMER or HHsearch (49). Similarly, we only considered matches with an E-value of less than 0.001. We used the FunMod modelling platform to build structural models. For those events where we failed to build a good model, we mapped them to the best structural matches (based either on the HHsearch or the HMMER result). We only analyse MXE events where both isoforms map to the same structural template. We aligned the isoforms with the structural template using MAFFT and extracted the variable residues between the two isoforms.

For those MXE events which we failed to annotate with structure information, we predicted if they were intrinsically disordered using IUPRED (50). We used the long-disorder option of IUPRED and residues with IUPRED score above 0.5 were considered disordered. We defined a splice isoform as intrinsically disordered if more than 50% of the residues were predicted to be disordered.

### Quantifying the residue changes using a physiochemical score

We compared the physiochemical properties of variable residues using the McLachlan physiochemical similarity matrix (31). A pair of amino acids was given a similarity score ranging from 0 to 6. A score of 0 indicates no similarity or a deletion. The score for a pair of identical amino acids is typically 5 or 6 (31). We summed up all the McLachlan similarity scores for a set of variable residues and divided by the total number of mapped variable residues to normalise the score.

### Analysis to determine if variable residues in the splice event are exposed to solvent

NACCESS (51) is a stand-alone program that calculates the relative accessible surface (rASA) of amino acids in a PDB structure. For each splice event, we calculated the rASA for all the corresponding variable residues of the two isoforms. Amino acid residues were considered to be exposed if the rASA value was above 10%. We compared the solvent exposure of the variable residues versus the solvent exposure of the non-variable residues. The solvent exposure was calculated as:

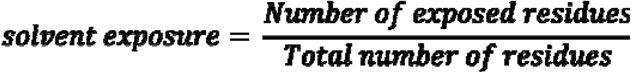

We also investigated random models to examine the solvent exposure for the splice region itself relative to background. For every MXE pair, we randomly selected from the same structure regions that have the same size as the splice region, 10,000 times, and determined the average solvent exposure of this region.

### Analysis to determine if variable residues in MXE regions are clustered in 3D and are close to functional sites

We investigated whether exposed variable residues in MXE events lie in the vicinity of functional residues. To reduce the noise and identify those variable residues likely to be having similar impacts, we clustered variable residues into structural clusters. For each pair of MXE events, we calculated the all-atom-versus-all-atom distances of the variable residue atoms of the structural representative and determ - - distance of each other (i.e. between the centres of the residues). We made sure there are at least three residues present in a cluster.

Subsequently, we calculated the distance from the structural clusters to known functional residues of the structural representative. We calculated the centre-of-mass of the clusters. The minimum distance between the centre-of-mass of

- to define if the structural cluster lies close to a functional site.

We also investigated random models to examine the proximity of randomly selected residue regions to functional sites, to evaluate statistical significance. For every MXE pair, we randomly selected structural clusters that have the same size as the largest MXE structural cluster, 10000 times, and determined the percentage of clusters that lie close to functional sites. This was done by randomly selecting a residue and then identifying the residues that are close (less than or equal to 2A) to the selected residue, and each other, until they made up the same volume as the largest MXE cluster. Then, the overall percentage for all random events was computed. This percentage is then compared with the actual MXE percentage. We used the z-score test to compute the statistical significance.

The functional sites considered were known experimentally characterised sites: catalytic residues taken from CSA (32), protein-protein interaction residues and protein-small molecule interaction residues (ligand binding) taken from IBIS (33). We also used our in-house predicted functional sites, FunSites (27), and predicted allosteric sites. For the prediction of FunSites, we only perform this analysis for FunFams that have a diverse set of relatives making it possible to distinguish conserved positions from variable positions. The Scorecons method (52) was used to calculate the sequence diversity of a FunFam alignment by generating a diversity of position score (DOPS score). FunFams with a DOPS score of above 70 are deemed sufficiently sequence diverse for analysis (i.e. having a lower probability of predicting false positives for Funsites (28)).

We also used the A-SITE allosteric site predictor (Dr Aurelio Moya Garcia, personal communication), based on the centrality of nodes (in a protein i.e. residue nodes) in terms of their capability to transfer signals through the protein, to predict allosteric sites in the protein, in order to determine whether variable residues lay on or close to allosteric sites. The program identifies residues with high betweenness centrality. The depletion of residues having high betweenness centrality values would be expected to interrupt the allosteric communication among regions of the protein that lie far apart. We defined the allosteric residues as the top 5 percentile residues (ranked by the A-SITE program using the betweenness centrality measure).

### Analysis to determine if variable regions in human MXE pairs are close to COSMIC cancer mutation residues

COSMIC is a database which collects somatic mutations from human cancer patients (34). Accumulation of these mutations may alter cellular functions and contribute to cancer (53). A number of studies have found that cancer mutation sites lie in the vicinity of functional residues (i.e. protein and ligand binding sites) (54, 55).

Recently, mutationally enriched functional families (MutFams) have been identified by in-house studies. MutFams are CATH functional families that have been found to be enriched in somatic missense cancer mutations reported in COSMIC. A total of 541 pan-cancer MutFams were identified and they indicate important cancer domain families such as P53 and PTEN (P. Ashford, personal communication). MutFam putative driver genes were found to have reasonable overlap with driver genes identified with genes from Cancer Gene Census and genes identified by other methods based on Pfam families. MutFam genes are enriched in survival and cell-motility cancer hallmark processes and additionally identify proteins with G1/S checkpoint function in DNA repair (P. Ashford, personal communication). Pan-cancer mutations clustered on representative domains in MutFams with available structural domains, have been found to be significantly closer to key functional sites than un-clustered cancer or germline disease mutations.

Therefore, we determined if MutFam cancer mutation residues lie close to MXE variable residues and functional residues. We mapped all the MutFam mutation residues onto the human MXE events. For those where we have structural information, we determined if the variable residue or residue clusters are close to MutFam mutation residues and functional sites using a distance cut-off of 4A□ (between any atoms in the residue pairs) or 6A□ (to the centre of the cluster).

## RESULTS

### Statistics of the MXE dataset

Table 1 summarises the number of MXE pairs and the number of genes identified for the five different species. Note that, the fly DSCAM genes have a total of 4 variable exon clusters giving 38,016 potential splicing isoforms. Because of the dominance of the DSCAM gene in the drosophila dataset we removed it from further analysis so as not to bias the results too heavily towards one protein.

**Table 1.** Number of MXEs identified in the different species analysed in this study.

| Species | Number of MXEs | Number of genes | Percentage Gene Ortholog Overlap Zebrafish MXE |
| --- | --- | --- | --- |
| Human | 143 | 125 | 14.5 |
| Fly | 259 | 84 | - |
| Mouse | 142 | 129 | 14.8 |
| Fugu fish | 237 | 184 | 14.5 |
| Zebra fish | 287 | 257 | - |

The MXE lengths span from 10 residues up to 400 residues, with a median size of 40 amino acids. The sequence identities of MXEs span from 25% up to 99% (See Figure 2).

**Figure 2.**
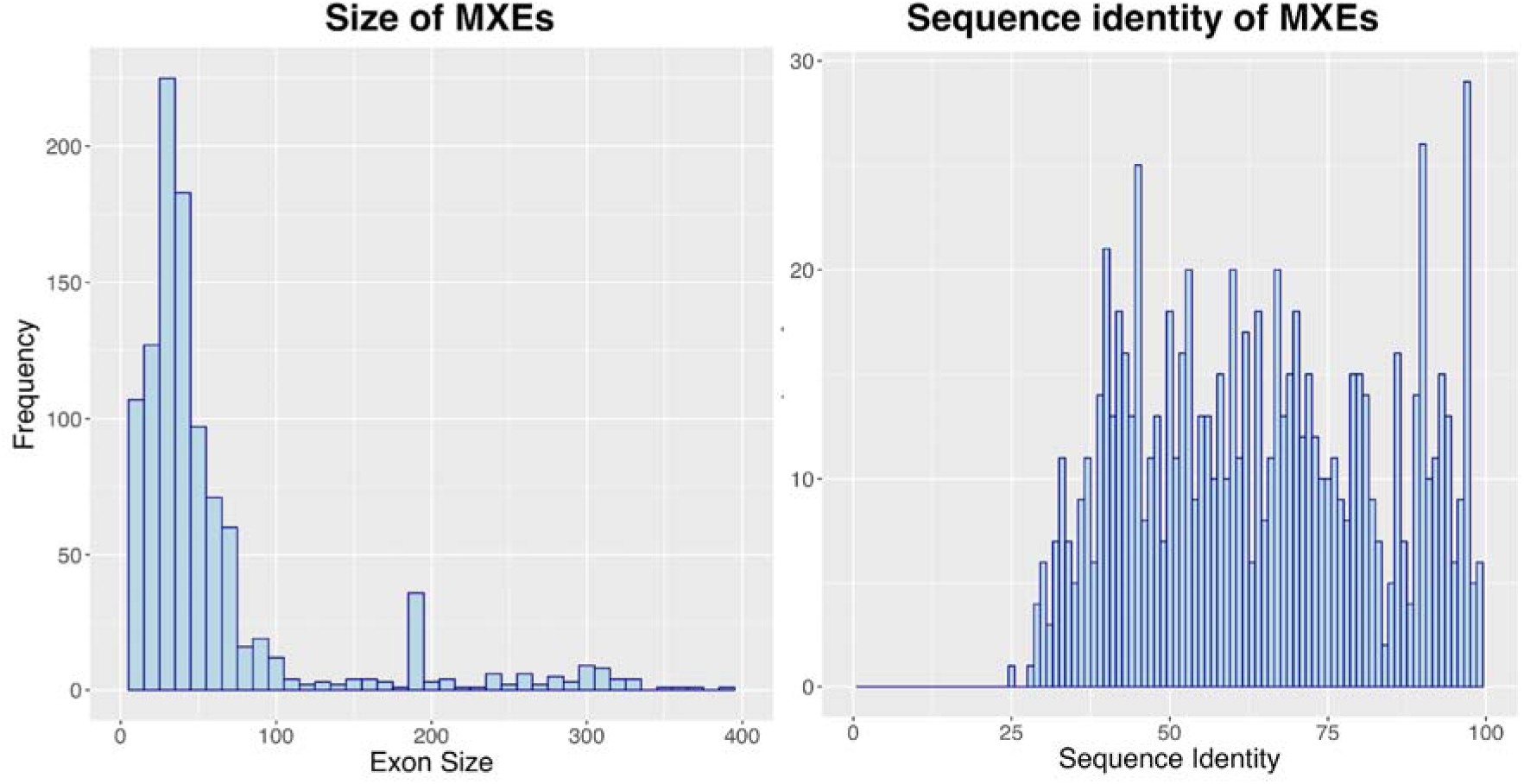
Size and sequence identity of MXEs. The exon size is given by the number of amino acid residues.

We also identified any orthologs for MXE genes between the species studied. We did this by using ensembl compara (including all orthology types 1 to 1 or 1 to many etc.) We then determined what percentage of genes in a given organism, containing an MXE event, had orthologous genes containing MXE events in other organisms. Human - Mouse MXE genes were found to have high overlap of 58%. However, this dropped to 30% for human compared to *zebra fish*. Interestingly when we considered the number of zebra fish MXE genes having an orthologous human MXE gene this coverage was only 15%. Furthermore, we found only 15% of zebra fish MXE genes orthologous to a Fugu gene with a MXE event (20% of Fugu MXE genes were identified as having an orthologous zebrafish MXE gene). This relatively low overlap between MXE events in the more distant species means that the strength of our conclusions will be reinforced by having applied a multi species approach to the analysis (i.e. since there is a high level of independence of gene sets (in terms of orthology groups) between species).

We used DAVID to identify the enriched biological annotations for these MXE genes relative to the whole genome background for each species. For all species, the MXE genes were enriched with membrane proteins (e.g. ion channels, synaptic vesicles and receptors) and enzymes (e.g. kinases, transferases, hydrolases and oxidoreductases), FDR level <0.01. (See Supplementary Figure 1 for MXE enriched terms).

We structurally annotated the splice isoforms with known protein structures and structural models built using the in-house FunMod modelling platform. For those events without a good model, we performed structural mapping to the representative structure of the respective FunFam by simply extracting the MAFFT alignment of the isoform against the sequence of the PDB structure. For those events that we failed to map to a FunFam, we performed structural modelling and structural mapping based on non-FunFam CATH domains, SCOPe domains or PDB chains. We also performed disorder analysis using IUPRED.

Figure 3 gives information on structural annotations for the sets of different splice isoforms. We managed to annotate 50% of the human isoforms with structural information. Less than 10% of the isoforms can be mapped to known structures. Less than 15% of the isoforms are disordered. We also determined the number of variable residues involved in MXE events. They usually involved less than 10 residues (See Figure 4).

**Figure 3.**
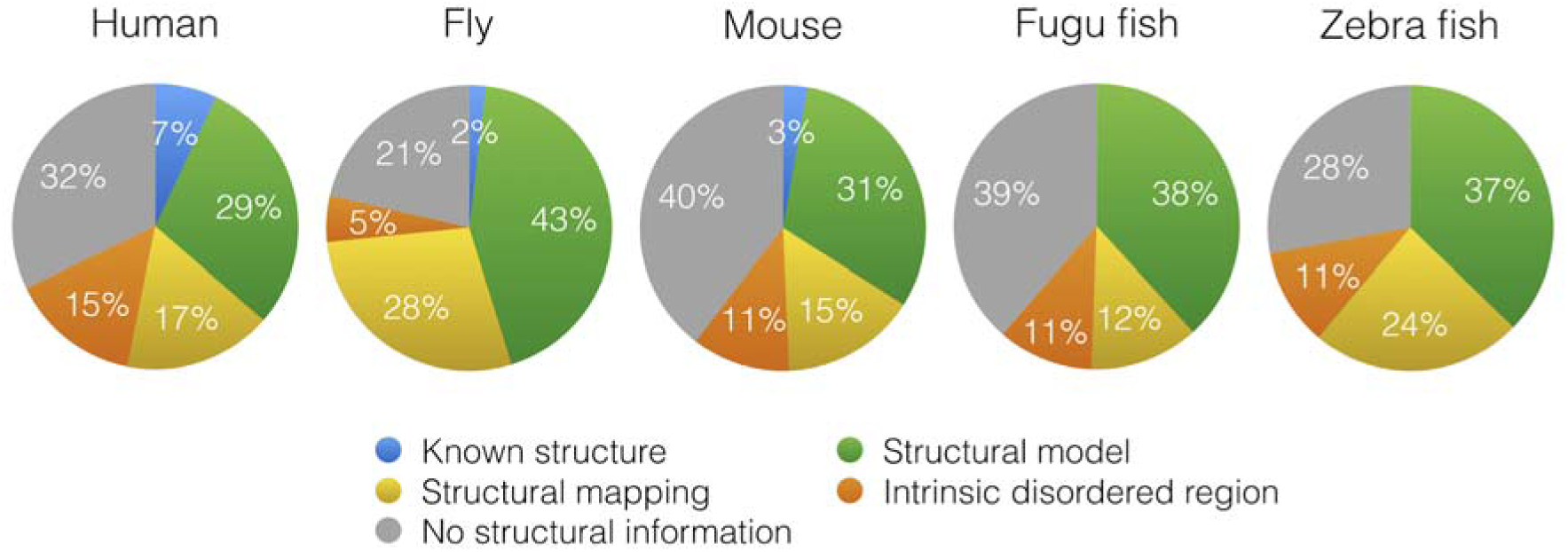
Splice isoforms with structural information.

**Figure 4.**
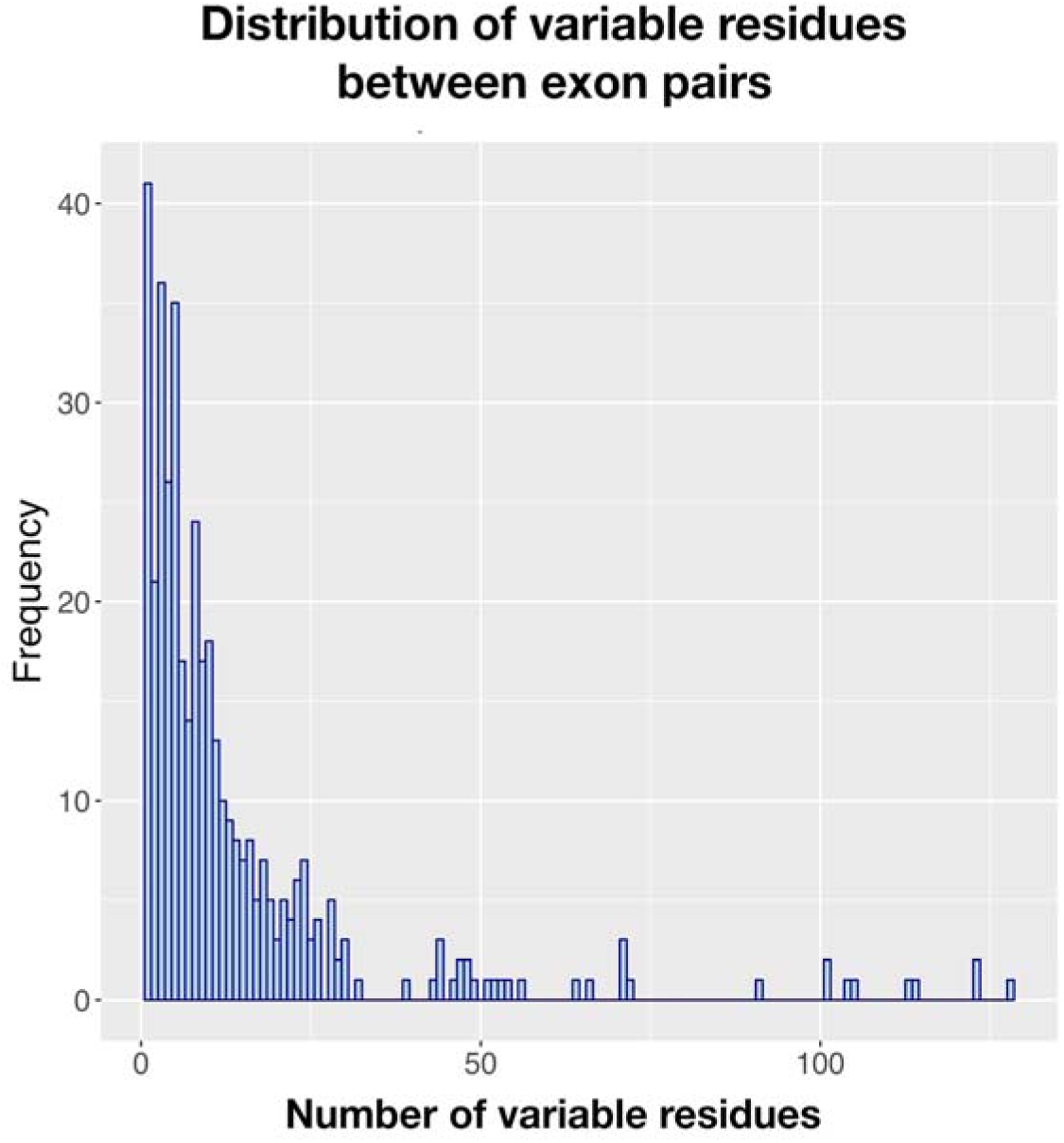
Number of variable residues between MXE exons in a pair

The FunFams mapped by the splice regions, from all species, are distributed among 115 CATH superfamilies. Figure 5 lists the names of CATH superfamilies common to all species. These are generally membrane proteins associated with cell-cell adhesion, signal transduction and molecule/ion transport.

**Figure 5.**
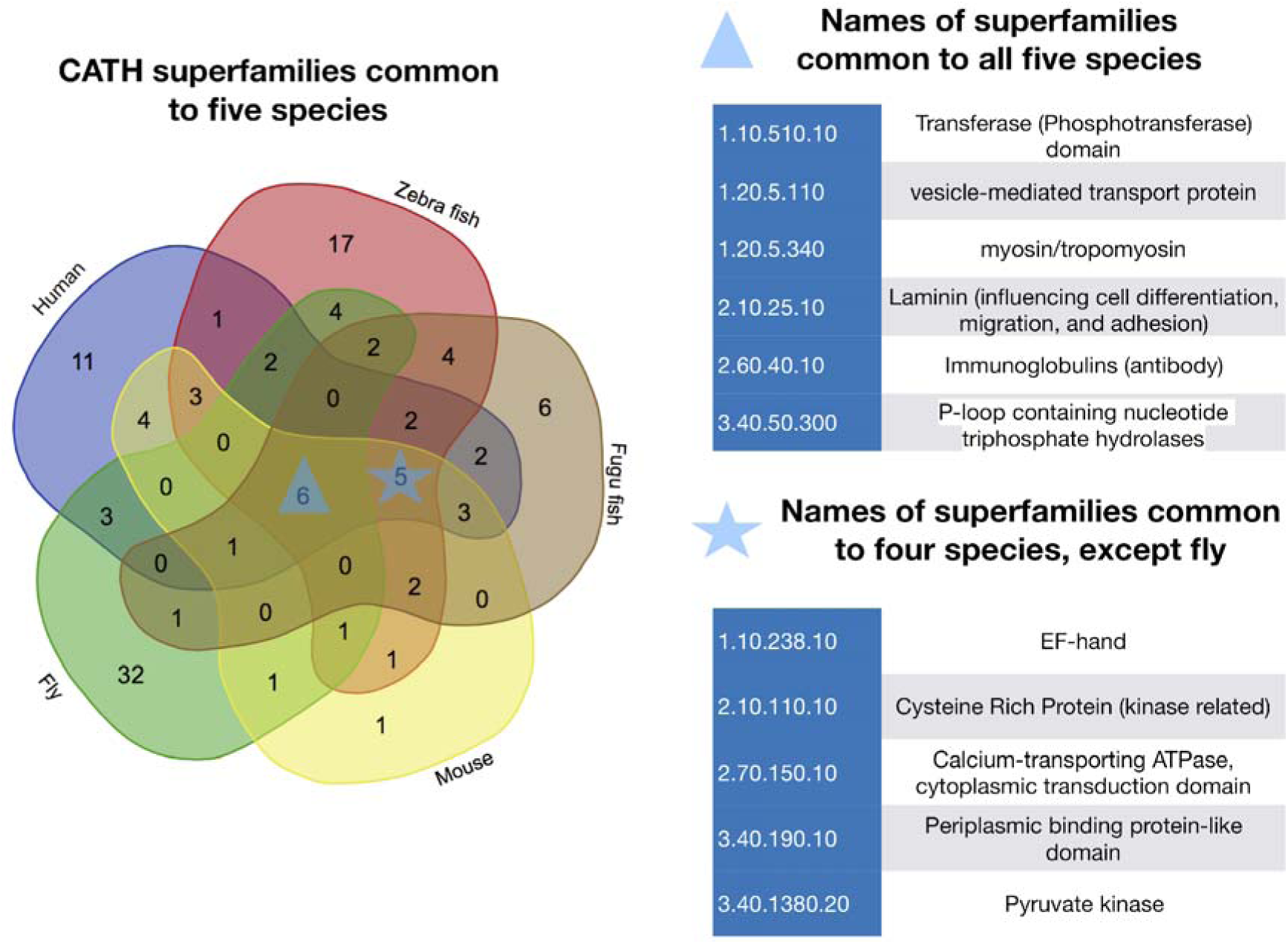
Number and names of CATH superfamilies common to all five species

### How often do MXE events involve a significant change in physiochemical properties of the variable residues?

We compared the physiochemical properties of the equivalent residues between the two isoforms using the McLachlan physiochemical matrix which captures chemical similarity between residues. Typically, McLachlan scores less than 2 for an amino acid change, are used to indicate a significant change in physiochemical properties. For all the datasets, more than 90% of the MXE variable residues have an average McLachlan score ≤2 (See Figure 6). These results indicate that in general the MXE events produce significant changes in physiochemical properties of the MXE regions. Such considerable changes suggest that MXE events are likely to cause functional shifts between protein isoforms where they are located within or close to functional sites in the protein.

**Figure 6.**
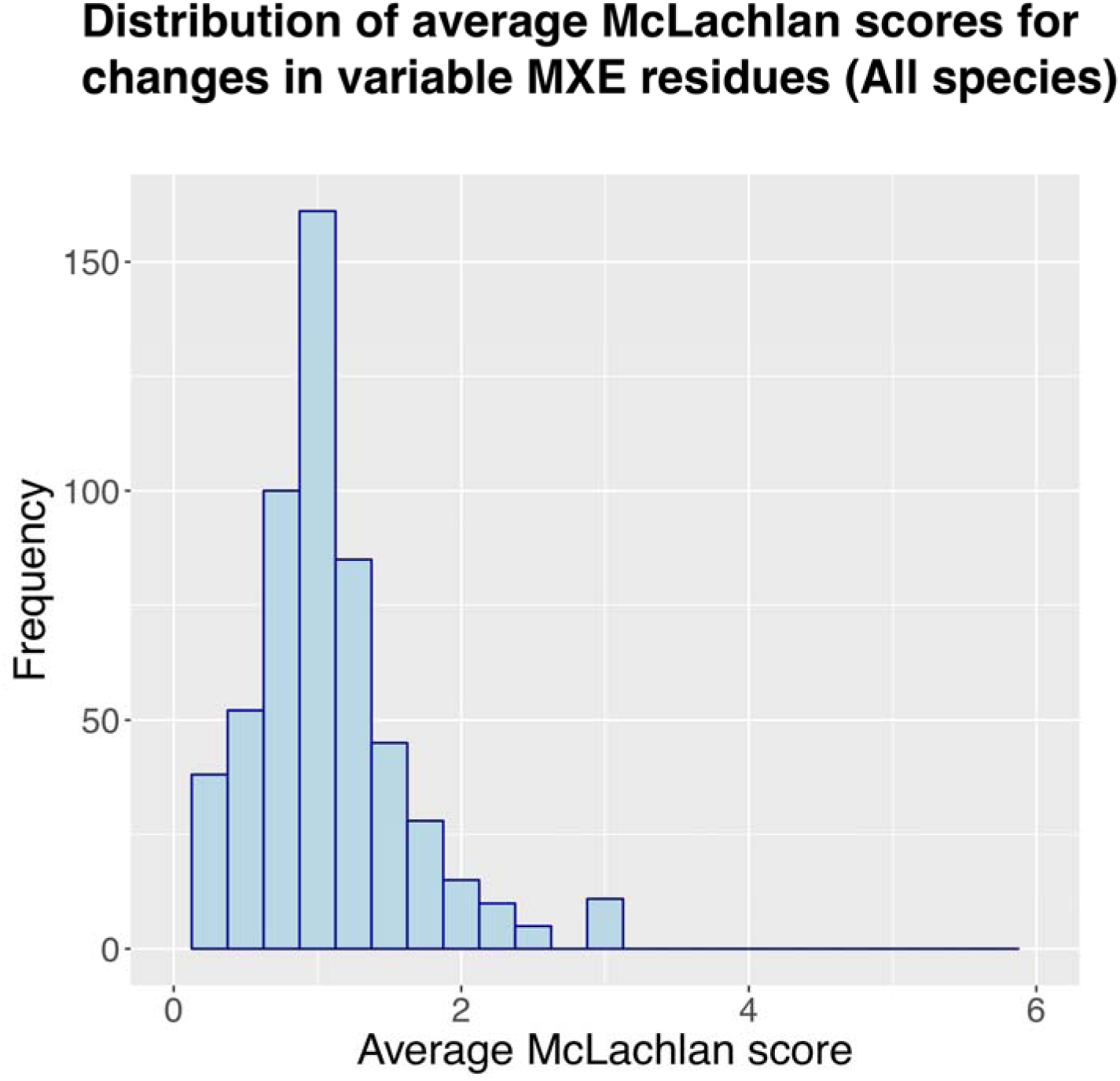
Distribution of average McLachlan scores for changes in variable MXE residues for all species.

### Analysing the frequency with which MXE events are exposed to solvent

To some extent, the function of a protein domain is largely dictated by the shape and nature of the residues on the protein surface. Enrichment of MXE events on surface residues would indicate an effect on function such as altered protein interactions etc. Because we had structural models of all our data, we were able to determine whether the MXE regions are more exposed than random (see methods). The percentage of exposed residues for a MXE region is significantly higher than random (Figure 7) (p-value <2.2e-16, Wilcoxon signed rank test). The level of the difference was striking and indicates that MXE events are strongly biased to altering residues with surface exposure. This picture was consistent across multiple organisms and indicates this is prominent and consistent feature of MXE splicing.

**Figure 7.**
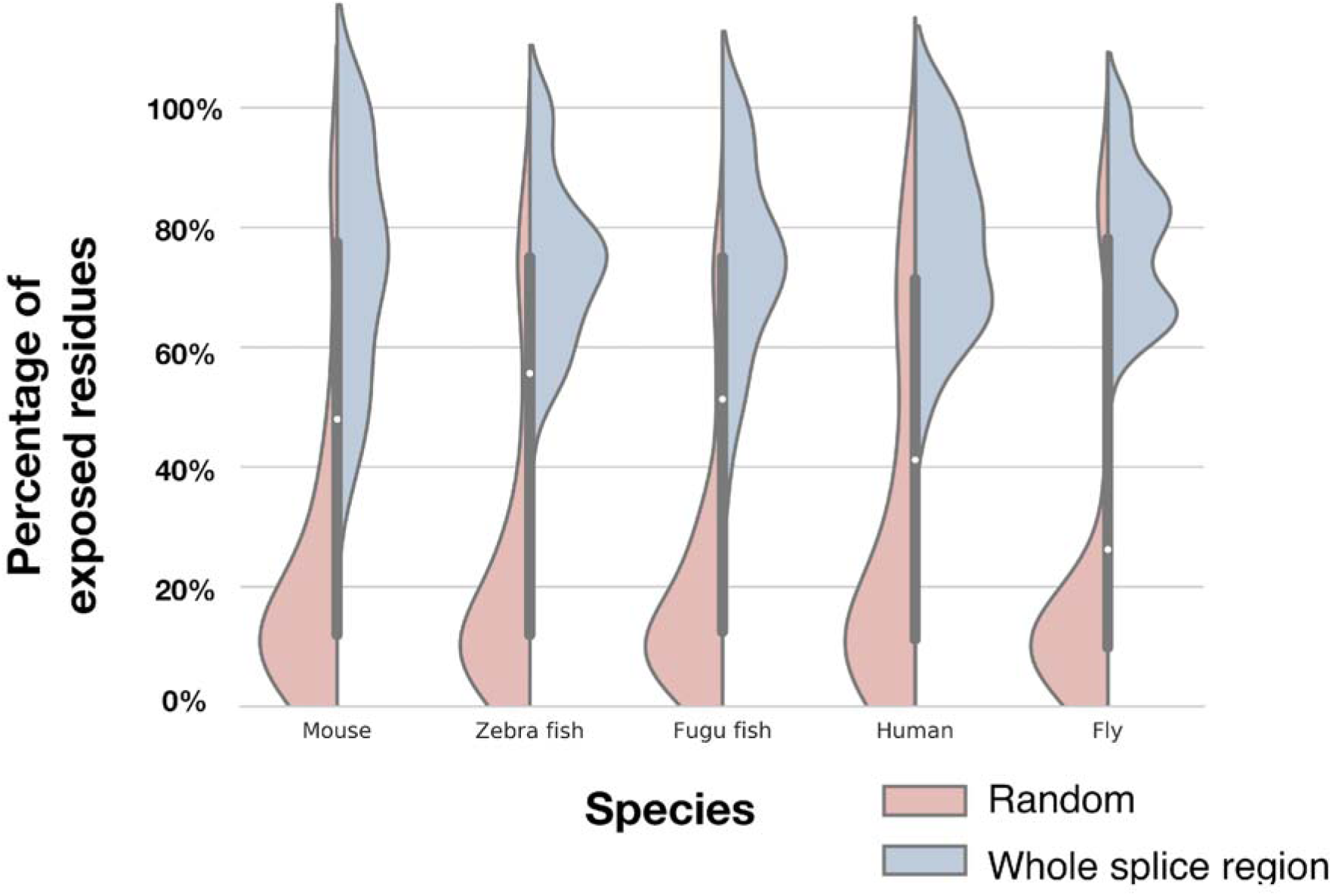
Proportion of exposed residues in the whole splice region versus a randomly selected region.

Next, we determined whether the variable residues involved in MXE events are more exposed to the solvent than the whole spliced region. Figure 8 compares the proportion of exposed variable MXE residues to the proportion of non-variable exposed MXE residues. Overall, we can observe that variable residues are more exposed than non-variable MXE residues. Again, the difference was highly significant (p-value <2.2e-16, Wilcoxon signed rank test). This result further confirms that the amino acid changes consistently alter surface exposed residues on the protein, much more than we would expect by random. We showed in the previous section that the McLachlan index changes are quite large for MXE events. The fact that MXE events tend to form stable proteins makes more sense if the changes are on the surface as we see here. Strong changes in amino acid properties at the surface would likely alter functions in binding partners or catalytic functions if close to interface or active site residues.

**Figure 8.**
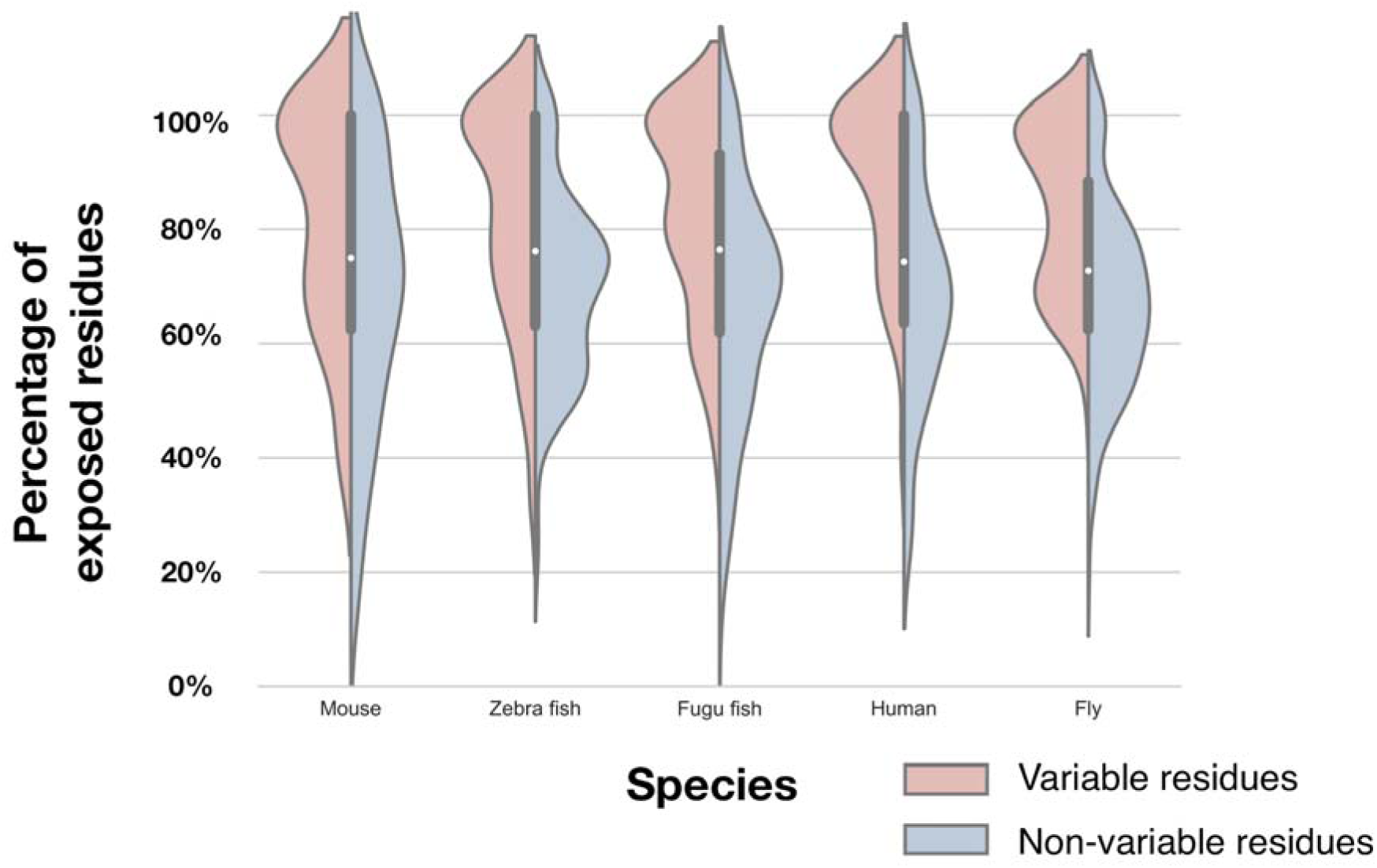
Proportion of exposed variable residues versus exposed non-variable residues in the whole of the splice region

### How often do clusters of variable residues in MXE events lie close to functional sites?

In the previous section, we demonstrated that the variable residues within the splice regions for splicing event pairs are more exposed to the solvent than the whole MXE region. Here, we investigate if these variable residues lie in the vicinity of functional residues. The known functional residues considered were catalytic residues, protein-protein interaction, protein-small molecules interaction sites. We also considered our in-house functional sites (FunSites) and allosteric sites.

In order to remove noise, we structurally clustered the variable residues. Clusters of variable resides were identified in over 70% of the MXE events. For those events that formed clusters, they usually formed 2-3 clusters (See Supplementary Figure 2).

Next, we determined whether structural clusters lie close to functional residues. For every pair of splice isoforms compared, we determined if the centre of mass of the MXE cluster was close to any functional sites (within 6A□). We calculated the minimum distance between the atoms of the residues involved. The results were compared with a random model - described in Methods.

Our results consistently showed that there is a significant tendency for clusters of MXE variable residues to lie close to protein-protein interaction, protein-small molecule, FunSites and allosteric sites for all species (Figure 9). We also performed the analysis without clustering the variable residues and obtained a similar result (see Supplementary Figure 3 for details). Through our structural modelling pipeline, integrated with functional site datasets, a clear picture emerges for a consistent role of MXE events modulating functional regions on domains. In support of previous work, our modelling suggests that unlike other forms of AS, the sequence variations are typically small and although chemically quite extreme (as judged by the McLachlan index), they are usually surface exposed and therefore unlikely to affect the ability to produce stable structures.

**Figure 9.**
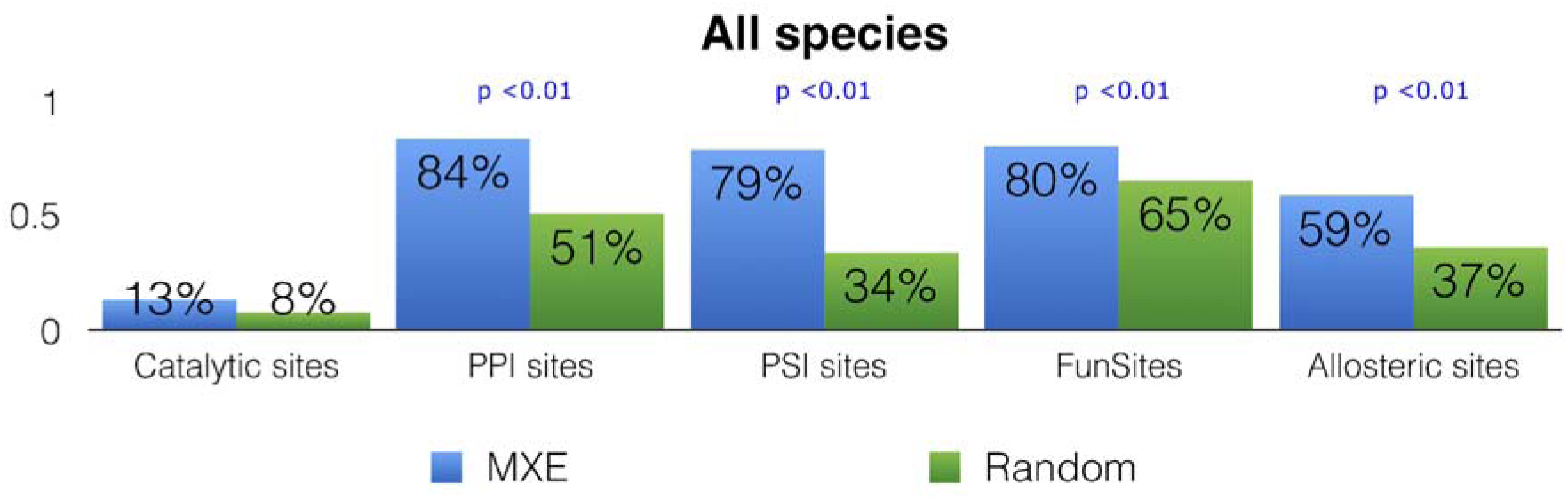
Proportion of variable region clusters in MXE events that lie close to functional sites compared to the number of random clusters that lie close to functional sites. NOTE: Plots for specific species can be found in the Supplementary Figure 4.

### How often do variable regions in MXE events lie close to putative cancer mutation sites?

A recent study by Hatje et al. (23) found that human MXEs were significantly enriched in pathogenic mutations. Furthermore, many studies have found that cancer mutation sites tend to lie in the vicinity of functional residues (i.e. protein and ligand binding sites) (54, 55). MutFams are CATH functional families found to be enriched in cancer mutations (see Methods). We therefore investigated whether MutFam cancer mutation residues coincide with both MXE variable residues and functional residues. Putative cancer mutations from MutFam were mapped to 153 of our human MXE events. We annotated 50 of these events with structures. We then determined if the variable residues/clusters in these MXE events were close to MutFam mutation residues. We found that 46 of the 50 MXE events had variable residues that were close to cancer mutations (statistically significant with p-value <0.01, compared with random models). We obtained the same results for both per-residue and per-cluster analyses. We checked if these MXE events were close to functional residues and found 40 out of 46 of the events were close to functional residues. A significant number of MutFam mutations are overlapping with MXE clusters which are close to functional sites, suggesting similarity in the mechanisms of any functional effects induced by alternative splicing and cancer mutations.

### Some examples of proteins displaying likely functional shifts following a MXE event

In this section, we present three examples that illustrate the impacts of MXE events on the function of the proteins.

#### Collapsin Response Mediator Protein (CRMP-PA versus CRMP-PB)

CRMP (FBgn0023023) is a dihydropyrimidinase enzyme (EC 3.5.2.2). It is predicted to be involved in biological processes such as centrosome localization, pyrimidine nucleobase biosynthetic process, and positive regulation of Notch signalling pathway. Currently, there is no experimental structure for the fly isoforms. The MXE exons can be mapped to FunFam (3.20.20.140.FF40287) which is populated with dihydropyrimidinase sequences. We modelled both of the isoforms using the same template (CATH domain 1gkpC02, also a dihydropyriminidase enzyme). The models generated have a GA341 score of 1 and normalised DOPE score of −0.71 suggesting they are of good quality. The structures share a high structural similarity (SSAP Score above 91) and sequence similarity (79%) (See Figure 10 for a structural superposition with the splice regions highlighted). We used the program Adaptive Poisson-Boltzmann Solver (56) to calculate surface potentials of the two isoforms. Over 80% of exposed splice region in CRMP-PA is positively charged. In contrast, the surface potential for CRMP-PB has a more negative charge that covers 60% of the exposed splice region (See Figure 10). We found out that there is a change in the zinc-binding residue 192, with a histidine in isoform CRMP-PA and a glycine in isoform CRMP-PB. This change is likely to disrupt the binding of the zinc molecule in the CRMP-PB isoform (See Supplementary Figure 5). Zinc molecules act as cofactors in dihydropyrimidinase enzymes. CRMP isoforms have been shown to have different tissue distributions and functions in fly (57).

**Figure 10.**
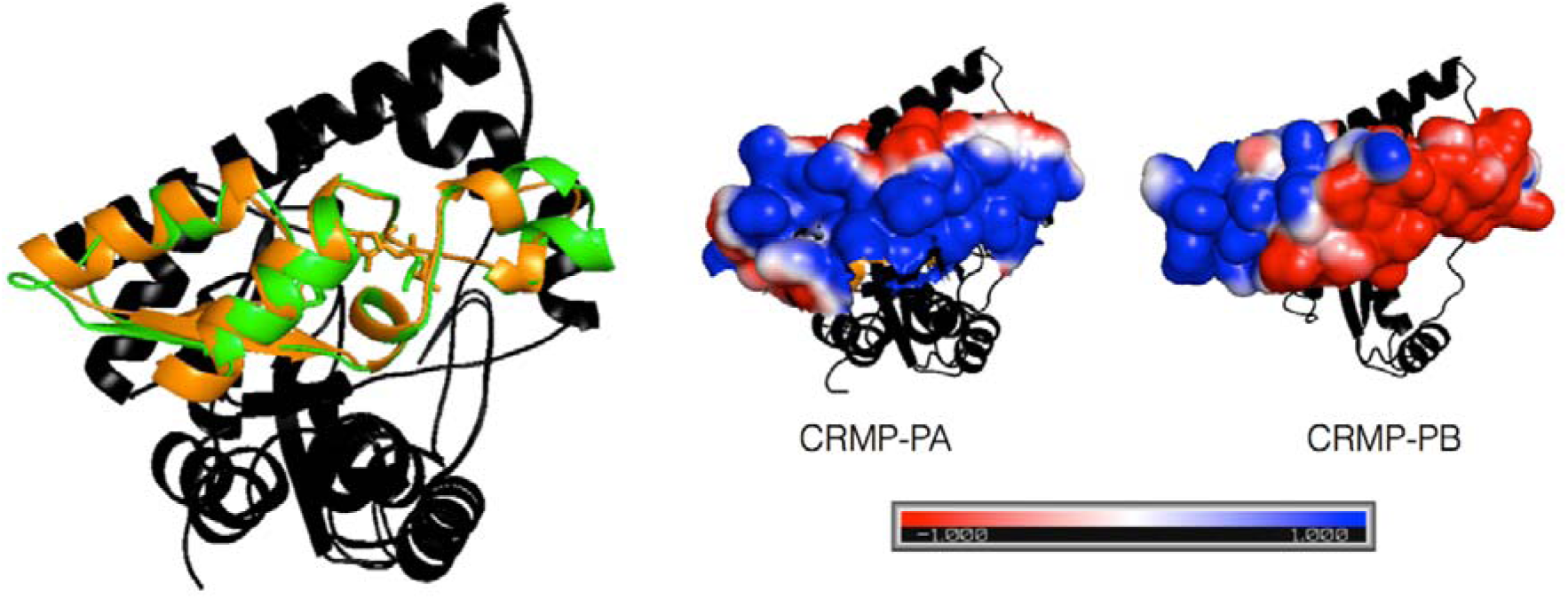
The splice regions of CRMP-PA (orange) and CRMP-PB (green) and surface potential of the splice regions

#### Pyruvate kinase (PKM1 versus PKM2)

The PKM gene codes for the dominant form pyruvate kinase, expressed in human tissue except liver and red blood cells. There are two isoforms associated with a MXE splicing event, known as PKM1 and PKM2. Most PKM expressing tissues, (irrespective of whether they are healthy or cancerous, embryonic or adult), express the M2 isoform of PK, while skeletal muscle and some other cell types that require a very high glycolytic flux, the constitutively active PKM1 isoform. Both the isoforms comprise 4 domains (See Supplementary Figure 6). The only difference between the isoforms is the MXE event located in the fourth domain (See Supplementary Figure 7). PKM1 is known to be a non-allosteric constitutively active tetramer (58). By contrast, PKM2 has been detected as an inactive monomer, a less active dimer, and a tetramer (inactive T-state and active R-state) which are regulated by metabolic intermediates and post-translational modifications (59, 60). Activity of the R-state tetramer PKM2 depends on an allosteric effector molecule fructose 1,6-bisphosphate (FBP), a situation that has been associated to the feed-forward regulation of glycolysis.

The regulation of pyruvate kinase activity plays an essential role in metabolism and is particularly important in the growth of cancer cells (60). PKM2 dimers and tetramers possess low and high levels of enzyme activity, respectively. Due to the key position of pyruvate kinase within glycolysis, the ability of PKM2 to switch between dimer and tetramer allows the tumour to modulate the flux of glycolysis, and with it energy production as the pyruvate kinase activity accounts for all glycolytic net ATP production. When the pyruvate kinase activity is high (ie when PKM2 is in the tetrameric state) and the flux of glycolysis is high, glycolytic ATP production is sufficient to meet the cellular demand; and most pyruvate formed from PKM2 is converted to lactate and excreted. When pyruvate kinase activity is low (ie when PKM2 is in the dimeric state), glycolytic ATP production is lower. Pyruvate is channelled at higher rates into the TCA cycle, in which it is fully oxidized to CO2, resulting in oxidative ATP production and the formation of TCA intermediates required for amino acid biosynthesis at higher rate. The increase in oxidative metabolism coincides with increased flux into the oxidative pentose phosphate pathway, producing higher levels of NADPH both required for anabolism and anti-oxidant enzymes (61, 62).

There are known structures available for both PKM1 (PDB id 3srf) and PKM2 (PDB id 1t5a) and domain 4 corresponds to CATH domains 3srfA01 and 1t5aA01. The overall sequence identity between the domains is high (87%) and the SSAP score between the domains is very good (SSAP score 95). See Supplementary Figure 6 for the structural superposition. It is known that PKM2 is activated by FBP but PKM1 is not.

Studies have shown that the FBP-binding residue 433 can be lysine, glutamate or threonine in different isoforms (63). There is a change in position 433 between PKM1 and PKM2, with a lysine in PKM2 and a glutamate in PKM1. Mutation of either threonine (yeast PK) or lysine (PKM2) to glutamate abolishes the FBP binding ability (64, 65). Mutation of glutamate in PKM1 to lysine enables the PKM1 mutant to bind FBP (66), presumably because FBP is negatively charged and thus a positively charged or neutral amino acid at this position would facilitate FBP binding whilst negatively charged amino acids would inhibit FBP binding and activation (63). Therefore, the presence of lysine in PKM2 facilitates the binding of FBP.

To form the tetramer, domain 4 of the PKM monomers interact (See Figure 11). The tetramer interface is located within the MXE region. We used the program Adaptive Poisson-Boltzmann Solver to calculate surface potentials of MXE regions (See Supplementary Figure 8). The MXE region for PKM1 has a higher surface complementary compared to PKM2 facilitating tetramer formation.

**Figure 11.**
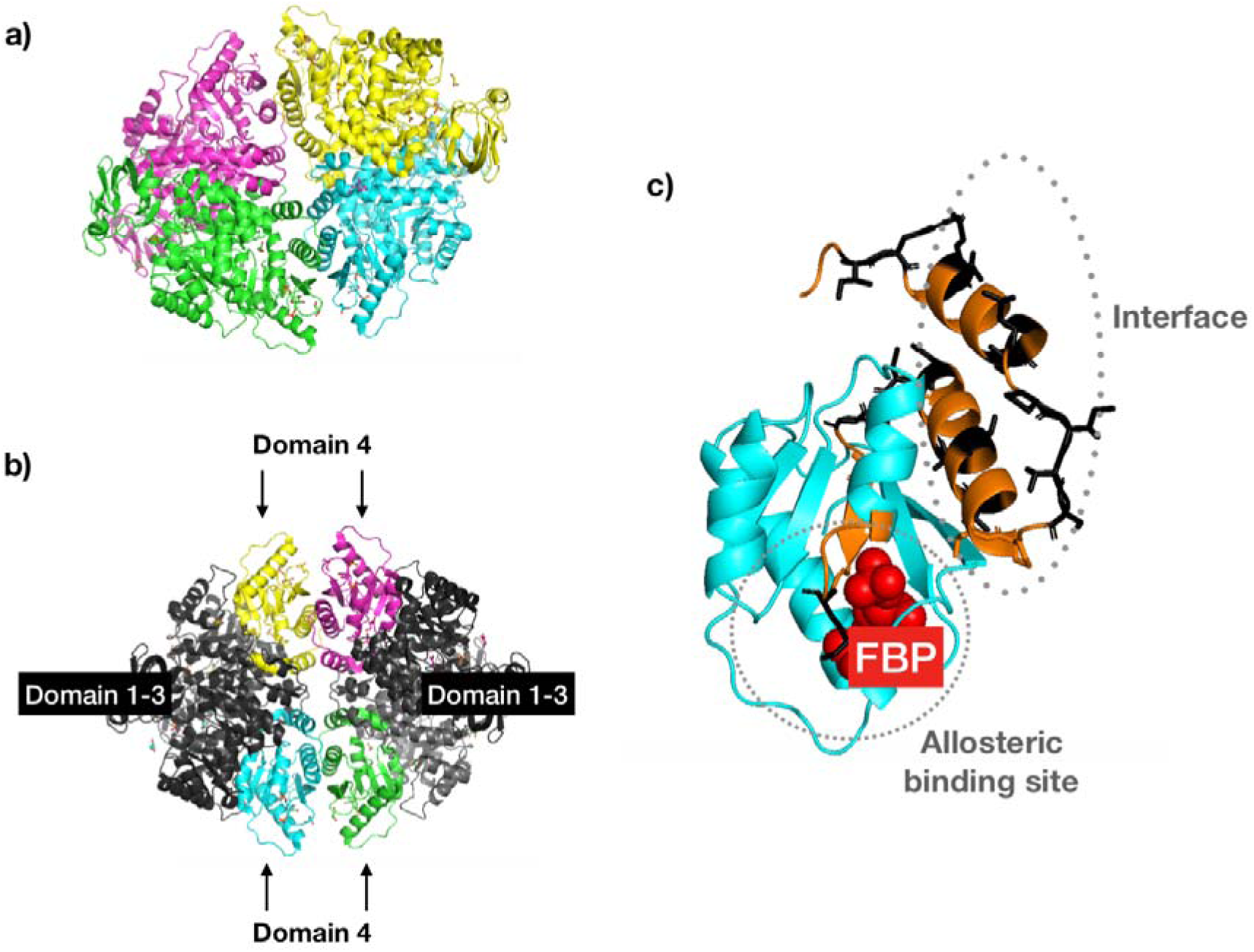
(a) The tetra-homomeric conformation of PKM. Each PKM monomer comprises four domains. The MXE event is located on domain 4. Domain 4 is highlighted in (b) and (c). The MXE region is coloured orange. Within the MXE region, the variable residues are coloured black. Variable residues are found near the FBP allosteric binding sites and in the interface region of the tetramer.

Although surfaces in the PKM2 interface are less complementary, studies have demonstrated that allosteric binding of FBP in PKM2 induces movement/rotations of the helices in the interface and the FBP activating loop that brings the tetramer binding residues into correct alignment for forming the tetramer (59, 67). Therefore, the lack of interface complementarity in PKM2 without FBP bound and the increase when FBP is bound facilitates the switch of PKM2 between dimer (without FBP) and tetramer (with FBP bound).

There are 15 variable residues within the MXE that lie within the tetramer interface formed by domain 4. We used mCSM (68) to assess the effect of the MXE variable residue changes on protein-protein affinity. A large number of the variable residues (14 out of the 15) changing from PKM1 to PKM2 were found to be destabilising to the tetramer complex (See Supplementary Figure 9). Similarly, 13 out of the 15 variable residues from PKM2 to PKM1 were found to be destabilising to the tetramer complex (See Supplementary Figure 10). We speculate this may be due to co-evolution of packing (i.e. for both PKM1 and PKM2, the interfaces have co-evolved to be complementary).

Both the MXE regions of PKM1/PKM2 were found to overlap with pan-cancer mutations in structural regions close to functional sites, reinforcing the idea that mutations in these regions impact on the function of the proteins.

#### Phosphofructokinase-platelet (PFKP-203 versus PFKP-205)

As demonstrated already in this paper, MXE events appear to be associated with functional switching events. However MXE events are also rare, occurring in only a small percentage of genes. As highlighted above glycolysis is of major importance and has disease associations with cancer. PKM has already been shown above to have MXE splicing with possible relevance to regulation, where the MXE event provides a mechanism for allosteric activation of this enzyme. Surprisingly, when we looked at the other major allosterically regulated enzyme in glycolysis, phosphofructokinase we also found a MXE event. PFK is a very tightly regulated enzyme (e.g. sensitive to ATP/ADP levels in the cell) and occupies the first “committed” step in glycolysis (effectively an irreversible reaction, that commits the sugar into the pathway) by catalysing the phosphorylation of fructose-6-phosphate to fructose-1,6-bisphosphate (FBP).

We find several variable residues between the MXE isoforms of PFKP including a change close to the FBP allosteric effector binding site (V->I) at residue 539 (See Figure 12). Although this is a conservative mutation, single amino acid changes from Valine to Isoleucine have previously been shown to change enzyme specificities (69). When we measured the change of FBP ligand binding affinity on mutating V to I at residue 539 with mCSM-lig (70), we found that the PFKP-205 isoform with the isoleucine residue binds FBP more efficiently (See Supplementary Figure 11) and we speculate this may affect the interaction of this isoform with FBP. Apart from being the product of the reaction, FBP is also a potent allosteric effector for PFKP (71). Binding of FBP induces conformational changes in PKKP that shifts the conformational equilibrium of PFKP to the activated R state (71). Given the role of PFK in disease and its key role in glycolysis the PFKP MXE could provide an interesting experimental study.

**Figure 12.**
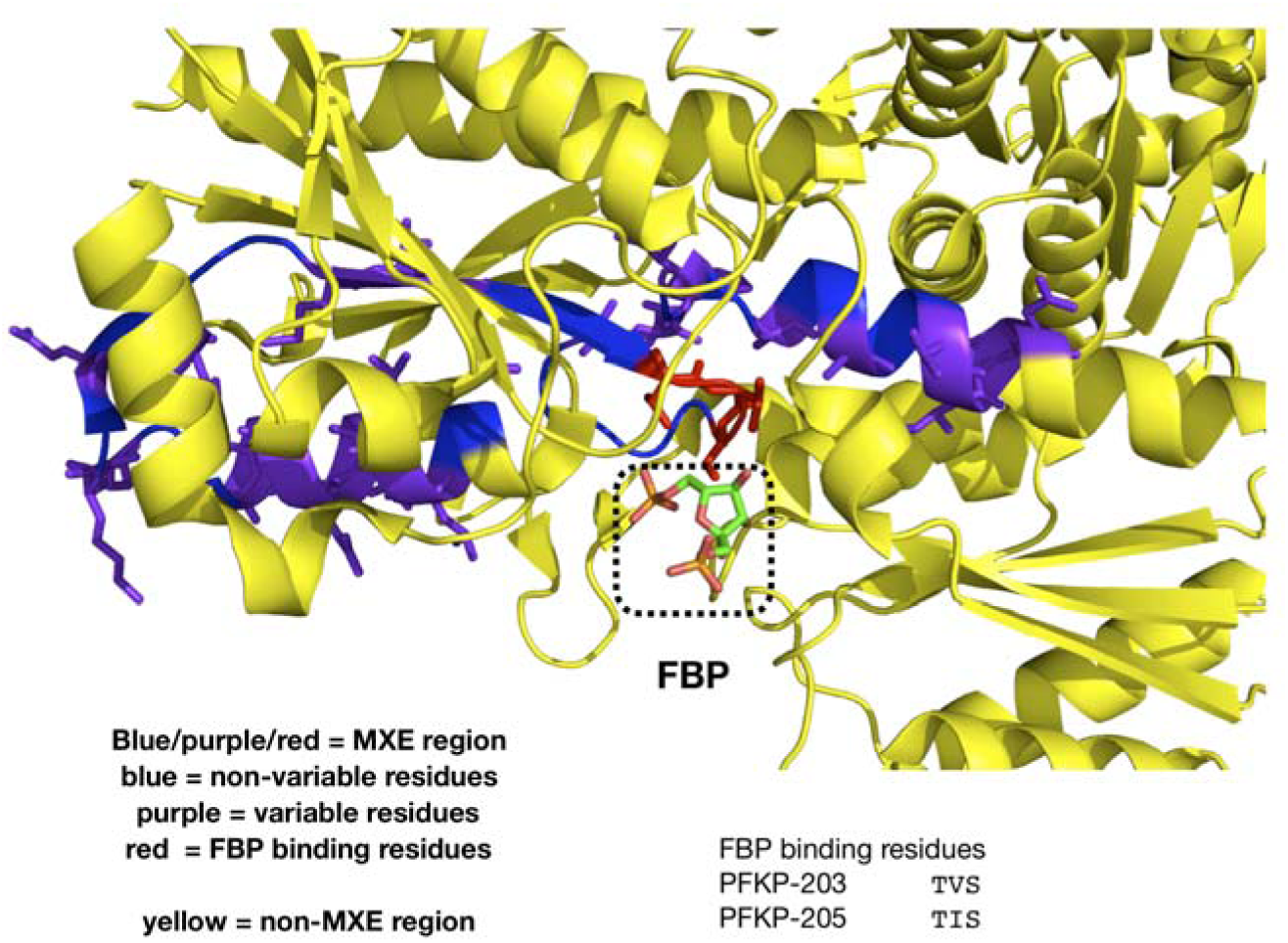
The structure of PFKP (PDB id 4xz2) with a mutually exclusive exon highlighted. Sequences come from (PFKP-203 (ENSP00000370517) and PFKP-205 (ENSP00000387871)).

## DISCUSSION

Mutually exclusive exons are characterised by coordinated splicing of exons such that only one of the two exons is retained, while the other is spliced out. These events are enriched in proteomics data, suggesting a functional role. Previous analyses by other groups have demonstrated that MXEs are evolutionary conserved among vertebrates (22, 23) and also demonstrated that these MXE regions are highly sequence similar and that MXE events do not usually disrupt the structural domain (14). However, to date, there are no reported studies in the literature on the likely functional consequences of MXE events, using structural models and five different species (human, fly, mouse, 2 fishes).

In this article, we identified the MXE events in human, mouse, fugu fish and zebrafish genomes using an in-house prediction protocol. The MXE lengths span from 10 residues up to 400 residues, with a median size of 40 amino acids. Most of these MXE genes are associated with membrane proteins (associated with cell-cell adhesion, signal transduction and molecule/ion transport) and enzymes (e.g. kinases, transferases, hydrolases and oxidoreductases). We annotated about 50% of the MXE events with structural information (i.e. known structures, structural models built using the FunMod modelling platform and structural mapping) and demonstrated that about 15% of the MXEs are disordered.

The fact that so many of the domains gave good structures (as judged by the structural quality scores such as N-DOPE) suggests that the MXE sequence changes do not affect the ability to form a stable protein. Although this has been considered likely to be the case from other independent studies, this is the first time this has been demonstrated using a large scale structural modelling analysis. Previous structural analyses have also shown that MXE events tend to be within structural domains (as opposed to being in e.g. intrinsically disordered regions). Our analysis shows that the MXE events although being within domains, have average surface exposure, far above what would be expected by random. We would expect this phenomena if the MXE switching events are associated with a change in function, as switching surface exposed residues is more likely to induce changes in protein functions (e.g. shifts in protein interfaces and binding pockets, etc.).

For those MXE events which can be mapped to a structure, they usually have less than 10 variable residues and they are likely to exhibit a significant change in their physiochemical property. We also demonstrated that variable residues are more exposed to the solvent than non-variable residues (within the same spliced region) and have a tendency to lie close to a known or predicted functional site (i.e. protein-protein interaction site, ligand binding site and allosteric site).

The fact that MXE regions are often in membrane proteins and or surface exposed identifies them as promising targets for druggable genes. Furthermore, there may a benefit in targeting a splice variant. For example, if a drug binds efficiently to its target but the inhibition of the protein causes general effects, leading to unwanted side effects, targeting a minor splice variant could be advantageous since its expression will be much more tissue or biological process specific. A splice variant specific drug could target a specific condition whilst leaving the more general function of the gene intact. A further advantage of MXE splice variants as drug targets is that, as we have demonstrated here, we can obtain reliable structural models for a large number of MXE events which could provide a good starting point for rational drug design / insecticides.

Recently, Hatje et al. predicted and validated more than 855 human MXEs, expanding the annotated human mutually exclusive exomes up to five fold (23). However, this dataset did not impose a sequence homology criterion in the MXE identification protocol. Furthermore, they did not perform comprehensive structural analyses as has been performed here. Hatje et al. found the human MXEs to be significantly enriched with pathogenic mutations (23). Here we expand this analysis to look at cancer mutations that occur in MutFams (CATH functional families found to be enriched in cancer mutations) and that fall in MXE regions. We find a significant proximity (in structure) between cancer mutations in these families and the MXE variable residues, suggesting a putative functional role for MXEs in human disease.

More generally we observed that MXE events are found in important functional classes of proteins such as key metabolic and signalling proteins. Why do some proteins expand their functional repertoire through MXE events and not others remains an unanswered question. There could be some high level of association between the genes that connects them, that has not yet been considered. For example, we might suspect that MXE functional expansion is somehow advantageous for genes expressed in certain tissues. We found a significant association for all human MXE genes to be associated with sensory tissue (FDR 1.6e-41: Fold enrichment=3.2) or nervous system (FDR =1.9e-29 Fold enrichment = 2.3) using TopAnat (72, 73). However, we did not find the same level of enrichment for *D.rerio* MXE genes which although they had a significant association with nervous system (FDR=1.5e-7) only had a weak fold enrichment (Fold enrichment = 1.17).

We found that MXE events are present in the two most highly regulated enzymes of glycolysis (PKM and FPK) and we show ways in which the MXE events can provide key regulatory mechanisms for these proteins. Again why some enzymes have developed MXE methods for altering how they are regulated or changing their functional specificities, rather than duplicating to give genes having separate functionalities, remains an open question.

Our study shows that the MXE regions tend not to be completely buried core residues of a fold, but rather regions with at least some solvent exposure, near to surface functional residues. MXE exons are ‘internal paralogues’ in that individual exons are duplicated and maintained, rather than whole genes. Although as we discussed MXE splicing is relatively rare, recent analysis has shown it is considerably more common than previously thought and that there are many examples of tandem arrays of duplicated MXE exons (23). Since we find that the MXE regions tend to be surface exposed, mutations in the MXE regions are less likely to disrupt core functional packing regions that are already highly optimised but instead produce changes that alter surface functional residues. Expression analysis shows that one MXE tends to have dominant expression (23) with other MXEs showing more specific tissue or developmental expression. The presence of a dominant isoform would allow the duplicated exon to diverge in sequence without loss of initial function. Our structural analysis shows that, mutations in the non-dominant (paralogous) MXE regions would be more likely to alter various surface functional processes (e.g. ligand binding). Once a novel function emerges in an MXE it may provide some novel advantageous function to a specific tissue/cell type.

Coupled with the fact that MXE splicing tends to be preserved at the proteomics level, our results suggest MXE events may have a number of important roles in cells generally.

## SUPPLEMENTARY DATA

Supplementary Data are available at NAR online.

## ACKNOWLEDGEMENT

The authors thank David Jones for fruitful discussions and Markus Ralser for his helpful comments on the PKM study.

## REFERENCES

1. Nilsen, T.W. and Graveley, B.R. (2010) Expansion of the eukaryotic proteome by alternative splicing. Nature, 463, 457–463.

2. Mauger, O. and Scheiffele, P. (2017) Beyond proteome diversity: alternative splicing as a regulator of neuronal transcript dynamics. Curr. Opin. Neurobiol., 45, 162–168.

3. Pohl, M., Bortfeldt, R.H., Grützmann, K. and Schuster, S. (2013) Alternative splicing of mutually exclusive exons—A review. Biosystems, 114, 31–38.

4. Tazi, J., Bakkour, N. and Stamm, S. (2009) Alternative splicing and disease. Biochim. Biophys. Acta - Mol. Basis Dis., 1792, 14–26.

5. Chen, J. and Weiss, W. a (2014) Alternative splicing in cancer: implications for biology and therapy. Oncogene, 34, 1–14.

6. Yang, X., Coulombe-Huntington, J., Kang, S., Sheynkman, G.M., Hao, T., Richardson, A., Sun, S., Yang, F., Shen, Y.A., Murray, R.R., et al. (2016) Widespread Expansion of Protein Interaction Capabilities by Alternative Splicing. Cell, 164, 805–817.

7. Light, S. and Elofsson, A. (2013) The impact of splicing on protein domain architecture. Curr. Opin. Struct. Biol., 23, 451–458.

8. Liu, S. and Altman, R.B. (2003) Large scale study of protein domain distribution in the context of alternative splicing. Nucleic Acids Res., 31, 4828–4835.

9. Buljan, M., Chalancon, G., Eustermann, S., Wagner, G.P., Fuxreiter, M., Bateman, A. and Babu, M.M. (2012) Tissue-specific splicing of disordered segments that embed binding motifs rewires protein interaction networks. Mol. Cell, 46, 871–883.

10. Ellis, J.D., Barrios-Rodiles, M., Çolak, R., Irimia, M., Kim, T., Calarco, J.A., Wang, X., Pan, Q., O’Hanlon, D. and Kim, P.M. (2012) Tissue-specific alternative splicing remodels protein-protein interaction networks. Mol. Cell, 46, 884–892.

11. Uhlén, M., Fagerberg, L., Hallström, B.M., Lindskog, C., Oksvold, P., Mardinoglu, A., Sivertsson, Å., Kampf, C., Sjöstedt, E., Asplund, A., et al. (2015) Tissue-based map of the human proteome. Science, 347, 1260419.

12. Tress, M.L., Martelli, P.L., Frankish, A., Reeves, G.A., Wesselink, J.J., Yeats, C., ĺsólfur Ólason, P., Albrecht, M., Hegyi, H., Giorgetti, A., et al. (2007) The implications of alternative splicing in the ENCODE protein complement. Proc. Natl. Acad. Sci., 104, 5495–5500.

13. Sánchez-Pla, A., Reverter, F., de Villa, M.C.R. and Comabella, M. (2012) Transcriptomics: mRNA and alternative splicing. J. Neuroimmunol., 248, 23–31.

14. Abascal, F., Ezkurdia, I., Rodriguez-Rivas, J., Rodriguez, J.M., del Pozo, A., Vázquez, J., Valencia, A. and Tress, M.L. (2015) Alternatively spliced homologous exons have ancient origins and are highly expressed at the protein level. PLoS Comput Biol, 11, e1004325.

15. Tress, M.L., Abascal, F. and Valencia, A. (2016) Alternative splicing may not be the key to proteome complexity. Trends Biochem. Sci., 42, 98–110.

16. Floor, S.N. and Doudna, J.A. (2016) Tunable protein synthesis by transcript isoforms in human cells. Elife, 5.

17. Weatheritt, R.J., Sterne-Weiler, T. and Blencowe, B.J. (2016) The ribosome-engaged landscape of alternative splicing. Nat. Struct. Mol. Biol., 23, 1–9.

18. Guttman, M., Russell, P., Ingolia, N.T., Weissman, J.S. and Lander, E.S. (2013) Ribosome profiling provides evidence that large noncoding RNAs do not encode proteins. Cell, 154, 240–251.

19. Tress, M.L., Abascal, F. and Valencia, A. (2017) Most Alternative Isoforms Are Not Functionally Important. Trends Biochem. Sci., 42, 408–410.

20. Pillmann, H., Hatje, K., Odronitz, F., Hammesfahr, B. and Kollmar, M. (2011) Predicting mutually exclusive spliced exons based on exon length, splice site and reading frame conservation, and exon sequence homology. BMC Bioinformatics, 12, 270.

21. Hatje, K. and Kollmar, M. (2014) Kassiopeia: a database and web application for the analysis of mutually exclusive exomes of eukaryotes. BMC Genomics, 15, 115.

22. Abascal, F., Tress, M.L. and Valencia, A. (2015) The evolutionary fate of alternatively spliced homologous exons after gene duplication. Genome Biol. Evol., 7, 1392–1403.

23. Hatje, K., Rahman, R., Vidal, R.O., Simm, D., Hammesfahr, B., Bansal, V., Rajput, A., Mickael, M.E., Sun, T., Bonn, S., et al. (2017) The landscape of human mutually exclusive splicing. Mol. Syst. Biol., 13, 959.

24. Chen, M., David, C.J. and Manley, J.L. (2012) Concentration-dependent control of pyruvate kinase M mutually exclusive splicing by hnRNP proteins. Nat. Struct. Mol. Biol., 19, 346–354.

25. Soom, M., Gessner, G., Heuer, H., Hoshi, T. and Heinemann, S.H. (2008) A mutually exclusive alternative exon of slo 1 codes for a neuronal BK channel with altered function. Channels, 2, 278–282.

26. Dawson, N.L., Sillitoe, I., Lees, J.G., Lam, S.D. and Orengo, C.A. (2017) CATH-Gene3D: Generation of the resource and its use in obtaining structural and functional annotations for protein sequences. In Methods in Molecular Biology. Vol. 1558, pp. 79–110.

27. Das, S., Lee, D., Sillitoe, I., Dawson, N.L., Lees, J.G. and Orengo, C.A. (2015) Functional classification of CATH superfamilies: a domain-based approach for protein function annotation. Bioinformatics, 31, 3460–3467.

28. Dessailly, B.H., Dawson, N.L., Mizuguchi, K. and Orengo, C.A. (2013) Functional site plasticity in domain superfamilies. Biochim. Biophys. Acta - Proteins Proteomics, 1834, 874–889.

29. Lam, S.D., Das, S., Sillitoe, I. and Orengo, C. (2017) An overview of comparative modelling and resources dedicated to large-scale modelling of genome sequences. Acta Crystallogr. Sect. D, 73, 628–640.

30. Lam, S.D., Dawson, N.L., Das, S., Sillitoe, I., Ashford, P., Lee, D., Lehtinen, S., Orengo, C.A. and Lees, J.G. (2016) Gene3D: expanding the utility of domain assignments. Nucleic Acids Res., 44, D404–D409.

31. McLachlan, A.D. (1972) Repeating sequences and gene duplication in proteins. J. Mol. Biol., 64, 417–437.

32. Furnham, N., Holliday, G.L., De Beer, T.A.P., Jacobsen, J.O.B., Pearson, W.R. and Thornton, J.M. (2014) The Catalytic Site Atlas 2.0: Cataloging catalytic sites and residues identified in enzymes. Nucleic Acids Res., 42, D485–D489.

33. Shoemaker, B.A., Zhang, D., Tyagi, M., Thangudu, R.R., Fong, J.H., Marchler-Bauer, A., Bryant, S.H., Madej, T. and Panchenko, A.R. (2012) IBIS (Inferred Biomolecular Interaction Server) reports, predicts and integrates multiple types of conserved interactions for proteins. Nucleic Acids Res., 40, D834–D840.

34. Forbes, S.A., Beare, D., Boutselakis, H., Bamford, S., Bindal, N., Tate, J., Cole, C.G., Ward, S., Dawson, E., Ponting, L., et al. (2016) COSMIC: somatic cancer genetics at high-resolution. Nucleic Acids Res., 45, D777–D783.

35. Gramates, L.S., Marygold, S.J., Dos Santos, G., Urbano, J.M., Antonazzo, G., Matthews, B.B., Rey, A.J., Tabone, C.J., Crosby, M.A., Emmert, D.B., et al. (2017) FlyBase at 25: Looking to the future. Nucleic Acids Res., 45, D663–D671.

36. Camacho, C., Coulouris, G., Avagyan, V., Ma, N., Papadopoulos, J., Bealer, K. and Madden, T.L. (2009) BLAST+: architecture and applications. BMC Bioinformatics, 10, 421.

37. Huang, D.W., Sherman, B.T. and Lempicki, R.A. (2009) Systematic and integrative analysis of large gene lists using DAVID bioinformatics resources. Nat. Protoc., 4, 44–57.

38. Jiang, Y., Oron, T.R., Clark, W.T., Bankapur, A.R., D’Andrea, D., Lepore, R., Funk, C.S., Kahanda, I., Verspoor, K.M., Ben-Hur, A., et al. (2016) An expanded evaluation of protein function prediction methods shows an improvement in accuracy. Genome Biol., 17, 1–17.

39. Dawson, N.L., Lewis, T.E., Das, S., Lees, J.G., Lee, D., Ashford, P., Orengo, C.A. and Sillitoe, I. (2017) CATH: an expanded resource to predict protein function through structure and sequence. Nucleic Acids Res., 45, D289–D295.

40. Eddy, S.R. (2011) Accelerated profile HMM searches. PLoS Comput. Biol., 7, e1002195.

41. Yeats, C., Redfern, O.C. and Orengo, C. (2010) A fast and automated solution for accurately resolving protein domain architectures. Bioinformatics, 26, 745–751.

42. Sali, A. and Blundell, T.L. (1993) Comparative protein modelling by satisfaction of spatial restraints. Protein Struct. by distance Anal., 64, 779–815.

43. Shen, M. and Sali, A. (2006) Statistical potential for assessment and prediction of protein structures. Protein Sci., 15, 2507–2524.

44. Melo, F., Sánchez, R. and Sali, A. (2002) Statistical potentials for fold assessment. Protein Sci., 11, 430–448.

45. Taylor, W.R. and Orengo, C. a (1989) Protein structure alignment. J. Mol. Biol., 208, 1–22.

46. Katoh, K. and Standley, D.M. (2013) MAFFT multiple sequence alignment software version 7: improvements in performance and usability. Mol. Biol. Evol., 30, 772–780.

47. Fox, N.K., Brenner, S.E. and Chandonia, J.-M. (2014) SCOPe: Structural Classification of Proteins—extended, integrating SCOP and ASTRAL data and classification of new structures. Nucleic Acids Res., 42, D304–D309.

48. Berman, H.M., Westbrook, J., Feng, Z., Gilliland, G., Bhat, T.N., Weissig, H., Shindyalov, I.N. and Bourne, P.E. (2000) The Protein Data Bank. Nucleic Acids Res., 28, 235–242.

49. Remmert, M., Biegert, A., Hauser, A. and Söding, J. (2012) HHblits: lightning-fast iterative protein sequence searching by HMM-HMM alignment. Nat. Methods, 9, 173–175.

50. Dosztányi, Z., Csizmók, V., Tompa, P. and Simon, I. (2005) The pairwise energy content estimated from amino acid composition discriminates between folded and intrinsically unstructured proteins. J. Mol. Biol., 347, 827–839.

51. Hubbard, S.J. and Thornton, J.M. (1993) Naccess. Comput. Program, Dep. Biochem. Mol. Biol. Univ. Coll. London, 2.

52. Valdar, W.S.J. (2002) Scoring residue conservation. Proteins Struct. Funct. Bioinforma., 48, 227–241.

53. Martincorena, I. and Campbell, P.J. (2015) Somatic mutation in cancer and normal cells. Science, 349, 1483–1489.

54. David, A. and Sternberg, M.J.E. (2015) The contribution of missense mutations in core and rim residues of protein--protein interfaces to human disease. J. Mol. Biol., 427, 2886–2898.

55. Yamada, K.D., Nishi, H., Nakata, J. and Kinoshita, K. (2016) Structural characterization of single nucleotide variants at ligand binding sites and enzyme active sites of human proteins. Biophys. physicobiology, 13, 157–163.

56. Baker, N.A., Sept, D., Joseph, S., Holst, M.J. and McCammon, J.A. (2001) Electrostatics of nanosystems: application to microtubules and the ribosome. Proc. Natl. Acad. Sci., 98, 10037–10041.

57. Morris, D.H., Dubnau, J., Park, J.H. and Rawls, J.M. (2012) Divergent functions through alternative splicing: The Drosophila CRMP gene in pyrimidine metabolism, brain, and behavior. Genetics, 191, 1227–1238.

58. Israelsen, W.J. and Vander Heiden, M.G. (2015) Pyruvate kinase: Function, regulation and role in cancer. Semin. Cell Dev. Biol., 43, 43–51.

59. Wang, P., Sun, C., Zhu, T. and Xu, Y. (2015) Structural insight into mechanisms for dynamic regulation of PKM2. Protein Cell, 6, 275–287.

60. Wong, N., Ojo, D., Yan, J. and Tang, D. (2015) PKM2 contributes to cancer metabolism. Cancer Lett., 356, 184–191.

61. Grüning, N.M. and Ralser, M. (2011) Cancer: Sacrifice for survival. Nature, 480, 190–191.

62. Grüning, N.M., Rinnerthaler, M., Bluemlein, K., Mülleder, M., Wamelink, M.M.C., Lehrach, H., Jakobs, C., Breitenbach, M. and Ralser, M. (2011) Pyruvate kinase triggers a metabolic feedback loop that controls redox metabolism in respiring cells. Cell Metab., 14, 415–427.

63. Dombrauckas, J.D., Santarsiero, B.D. and Mesecar, A.D. (2005) Structural Basis for Tumor Pyruvate Kinase M2 Allosteric Regulation and Catalysis. Biochemistry, 44, 9417–9429.

64. Bond, C.J., Jurica, M.S., Mesecar, A. and Stoddard, B.L. (2000) Determinants of allosteric activation of yeast pyruvate kinase and identification of novel effectors using computational screening. Biochemistry, 39, 15333–15343.

65. Lyssiotis, C.A., Anastasiou, D., Locasale, J.W., Vander Heiden, M.G., Christofk, H.R. and Cantley, L.C. (2012) Cellular control mechanisms that regulate pyruvate kinase M2 activity and promote cancer growth. Biomed. Res., 23, 213–217.

66. Ikeda, Y., Taniguchi, N. and Noguchi, T. (2000) Dominant negative role of the glutamic acid residue conserved in the pyruvate kinase M1 isozyme in the heterotropic allosteric effect involving Fructose-1, 6-bisphosphate. J. Biol. Chem., 275, 9150–9156.

67. Morgan, H.P., O’Reilly, F.J., Wear, M.A., O’Neill, J.R., Fothergill-Gilmore, L.A., Hupp, T. and Walkinshaw, M.D. (2013) M2 pyruvate kinase provides a mechanism for nutrient sensing and regulation of cell proliferation. Proc. Natl. Acad. Sci., 110, 5881–5886.

68. Pires, D.E. V, Ascher, D.B. and Blundell, T.L. (2014) mCSM: predicting the effects of mutations in proteins using graph-based signatures. Bioinformatics, 30, 335–342.

69. Yuan, X., Yin, P., Hao, Q., Yan, C., Wang, J. and Yan, N. (2010) Single amino acid alteration between valine and isoleucine determines the distinct pyrabactin selectivity by PYL1 and PYL2. J. Biol. Chem., 285, 28953–28958.

70. Pires, D.E.V., Blundell, T.L. and Ascher, D.B. (2016) MCSM-lig: Quantifying the effects of mutations on protein-small molecule affinity in genetic disease and emergence of drug resistance. Sci. Rep., 6, 29575.

71. Kloos, M., Brüser, A., Kirchberger, J., Schöneberg, T. and Sträter, N. (2015) Crystal structure of human platelet phosphofructokinase-1 locked in an activated conformation. Biochem. J., 469, 421–432.

72. Bastian, F., Parmentier, G., Roux, J., Moretti, S., Laudet, V. and Robinson-Rechavi, M. (2008) Bgee: Integrating and comparing heterogeneous transcriptome data among species. In Lecture Notes in Computer Science (including subseries Lecture Notes in Artificial Intelligence and Lecture Notes in Bioinformatics).Vol. 5109 LNBI, pp. 124–131.

73. Komljenovic, A., Roux, J., Robinson-Rechavi, M. and Bastian, F.B. (2016) BgeeDB, an R package for retrieval of curated expression datasets and for gene list expression localization enrichment tests. F1000Research, 5, 2748.

